# Vaccine-linked chemotherapy with a low dose of benznidazole plus a bivalent recombinant protein vaccine prevents the development of cardiac fibrosis caused by *Trypanosoma cruzi* in BALB/c mice

**DOI:** 10.1101/2022.02.16.480638

**Authors:** Victor Manuel Dzul-Huchim, Maria Jesus Ramirez-Sierra, Pedro Pablo Martinez-Vega, Miguel Enrique Rosado-Vallado, Victor Ermilo Arana-Argaez, Jaime Ortega-Lopez, Fabian Gusovsky, Eric Dumonteil, Julio Vladimir Cruz-Chan, Peter Hotez, María Elena Bottazzi, Liliana Estefania Villanueva-Lizama

## Abstract

**Background:** Chagas disease (CD) is caused by *Trypanosoma cruzi* and affects 6-7 million people worldwide. Approximately 30% of chronic patients develop chronic chagasic cardiomyopathy (CCC) after decades. Benznidazole (BNZ), one of the first-line chemotherapy approved for CD, induces toxicity and fails to halt the progression of CCC in chronic patients. The recombinant parasite-derived antigens, including Tc24, Tc24-C4, TSA-1, and TSA-1-C4 with Toll-like receptor 4 (TLR-4) agonist-adjuvants reduce cardiac parasite burdens, heart inflammation, and fibrosis, leading us to envision their use as immunotherapy together with BNZ. Given genetic immunization (DNA vaccines) encoding Tc24 and TSA-1 induce protective immunity in mice and dogs, we propose that immunization with the corresponding recombinant proteins offers an alternative and feasible strategy to develop these antigens as a bivalent human vaccine. We hypothesized that a low dose of BNZ in combination with a therapeutic vaccine (TSA-1-C4 and Tc24-C4 antigens formulated with a synthetic TLR-4 agonist-adjuvant, E6020-SE) could provide antigen-specific T cell immunity and prevent cardiac fibrosis progression.

**Methodology/ Principal findings:** We evaluated the therapeutic vaccine candidate plus BNZ (25 mg/kg/day/7 days) given at days 72 and 79 post-infection (p.i) (early chronic phase). Fibrosis, inflammation, and parasite burden were quantified in heart tissue at day 200 p.i. (late chronic phase). Further, spleen cells were collected to evaluate antigen-specific CD4^+^ and CD8^+^ T cell immune responses, using flow cytometry. We found that vaccine-linked BNZ treated mice had lower cardiac fibrosis compared to the infected untreated control group. Moreover, cells from mice that received the immunotherapy had higher stimulation index of antigen-specific CD8^+^Perforin^+^ T cells as well as antigen-specific central memory T cells compared to infected untreated control.

**Conclusions:** Our results suggest that the bivalent immunotherapy together with BNZ treatment protects mice against cardiac fibrosis and activates strong CD8^+^ T cell responses by *in vitro* restimulation, evidencing the induction of a long-lasting *T. cruzi*-immunity.

**Author summary:** Chagas disease (CD) is a neglected tropical disease caused by the parasite *Trypanosoma cruzi,* transmitted through contact with infected feces of vectors bugs. CD can induce cardiac abnormalities including the development of fibrosis and eventually death. Benznidazole (BNZ) is the first-line drug approved against CD, however, its toxicity and lack of efficacy in the chronic phase have limited its use. Previous studies have demonstrated the feasibility of reducing doses of BNZ given in combination with therapeutic vaccines during the acute phase of CD, which increases its tolerability and reduces adverse side effects. Considering that patients are often diagnosed until prolonged stages of the disease, its necessary to evaluate therapies given in the chronic phase of CD. In this study, we evaluated a vaccine formulated with the recombinant *T. cruzi*-antigens TSA-1-C4 and Tc24-C4 and the adjuvant E6020-SE in combination with a low dose of BNZ given during the chronic phase of *T. cruzi*-infection using a murine model. The authors found that the combination therapy protects mice against cardiac fibrosis, allow the activation of a CD8^+^ T cell response and induce a prolonged memory response against *T. cruzi*. This study supports the development of the vaccine-linked chemotherapy approach in order to prevent *T. cruzi* chronic infection.

## Introduction

Chagas disease (CD) is caused by the protozoan parasite *Trypanosoma cruzi* (*T. cruzi*) transmitted mainly through contact with infected feces of hematophagous triatomine bugs. CD affects approximately 6.5 million people worldwide and is a major public health problem in Latin America (1). Moreover, CD is emerging in non-endemic regions due to human migration, political and socioeconomic instability, climate change and other factors (2). CD has two major clinical stages. The first is the acute infection and is characterized by high levels of parasites in peripheral blood; where individuals are mostly asymptomatic but can present non-specific febrile illness, which typically resolves within 4–8 weeks (3, 4). In the chronic phase, where approximately 20-30% of individuals develop chronic chagasic cardiomyopathy (CCC) years to decades after the initial infection, and some develop pathologies such as megaesophagus and megacolon (5, 6). The pathogenesis of CCC is due to *T. cruzi* persistence in the heart that drives chronic inflammation and fibrosis leading to abnormalities of the conduction system, e.g., right bundle branch block (RBBB), arrhythmias, tachycardia and subsequently death by heart failure (4,7,8). Benznidazole (BNZ), is one of the first-line chemotherapies used for CD treatment, however, it is associated with toxic side effects, has a poor efficacy in patients with chronic infection and they require long treatment, increasing the risk of drug resistance (9). Furthermore, therapy with BNZ does not reduce cardiac clinical deterioration through 5 years of follow-up in CCC patients (10). Conversely, other studies have reported that BNZ treatment is associated with a reduction in heart disease progression, suggesting that more trials focused on BNZ should be performed (11). The contrasting results between these studies may be explained by several factors as the number of individuals evaluated in each study, differences of geographical area where individuals are from, variation of parasite strains causing the infection, rates of loss to follow-up, and the diversity of clinical status of enrolled patients

The immune response required to reduce parasite dissemination, prevent cardiac lesions, and ensure host survival against *T. cruzi*, is still under investigation. The development and use of therapeutic vaccines are an alternative approach against CD. Its purpose is to stimulate the immune response from the host against *T. cruzi*-infection. Several therapeutic vaccine candidates have been evaluated exploring a vast of delivery systems (plasmids, adenoviruses, peptides and recombinant proteins) and adjuvants (12–16). Overall, in animal models of *T. cruzi* experimental infection, either T helper (Th)-1 or Th1/Th2 balanced and Th-17 immune responses are required to achieve parasite control (17–19) with evidence for the importance of IFNγ and CD8^+^ T cells (20, 21). A protective immunity is mediated by CD8^+^ cytolytic T cells (CTL), which release cytotoxic granules (perforin and granzymes) (21, 22). Perforin is a pore forming cytolytic protein, that allows the transition of granzymes, a group of serine proteases, which induce programmed cell death in *T. cruzi* infected cells (23, 24). In addition, CD8^+^ T cells modulate the immune response through the secretion of pro-inflammatory cytokines, such as IFNγ, IL-12 and tumor necrosis factor alpha (TNF-*α*) (25–27).

Our program has been examining the effects of two major recombinant protein antigens, together with Toll-like receptor 4 (TLR-4) agonist adjuvants. These antigens include a flagellar calcium-binding protein, Tc24, or a genetically re-engineered Tc24 antigen with cysteine modifications to prevent aggregation, known as Tc24-C4 (28, 29), and a trypomastigote surface antigen known as TSA-1 (30). Hence, both proteins are being produced under current good manufacturing practices (cGMP) as potential vaccines. Studies performed in acute *T. cruzi*-infection in mice indicate that both Tc24 and TSA-1 recombinant proteins drive a Th1 or balanced Th1/Th2 immunity in achieving therapeutic effects, with an emphasis on their role in reducing parasite persistence in the heart, and the associated with fibrosis and inflammation (15–17,31,32). To date, most of the studies in mice have focused on immunizations using a single recombinant protein antigen, nevertheless our earlier work using plasmid DNA immunization evaluating bivalent vaccines with both TSA-1 and Tc24 (33), demonstrated that there is a beneficial effect to use bivalent vaccines when are administered in mice during acute infection.

In addition, studies focused on evaluating the phenotype of memory T cells are important for vaccine development; identification of a vaccine that can effectively induce lasting memory-response is expected to prevent infection. According to the model proposed by Lanzavecchia and Sallusto (34), based on the expression of receptors required for lymph node homing, memory T cells are often classified as central memory (T_CM_), which can migrate to inflamed peripheral tissues and display immediate effector function, or effector memory (T_EM_) which remain in secondary lymphoid organs, have little or no effector function, but readily proliferate and differentiate to effector cells. Therefore, therapeutic vaccination strategies have focused on promoting antigen-specific CD4^+^ and CD8^+^ memory T cells during the chronic phase of *T. cruzi-*infection (35–38), which is also when most Chagasic patients seek treatment.

On the other hand, BNZ dosage and treatment regimens have been controversial in recent years. To assess the feasibility of minimizing the toxic side-effects, several trials have been performed to test low doses of BNZ given alone or in combination with other drugs or antigens (39–41). Accordingly, combined regimens improve the trypanocide activity and attenuate the BNZ toxicity, thus, these studies support the evaluation of an immunotherapy based on a reduced low-dose of BNZ linked to a vaccine formulated with the recombinant proteins TSA-1-C4 and Tc24-C4. Recently studies have demonstrated the efficacy of the recombinant Tc24-C4 antigen in combination with a low dose of BNZ administered during *T. cruzi* acute infection in a murine model resulting in increased antigen-specific CD8^+^ and IFNγ-producing CD4^+^ T cells populations, as well as in increased cytokines related to Th17 immune responses (42).

Although the bivalent vaccine has shown immunogenicity and protection in acute murine models, it needs to be evaluated in pre-clinical models of *T. cruzi* chronic infection. Hence, our group previously performed a pilot study in order to evaluate the progression of *T. cruzi* chronic infection in BALB/c mice based on the parasitemia and cardiac clinical manifestations evaluated using electrocardiograms (taken every 35 days) and echocardiograms (at 210 dpi). Those results suggested that days 72 and 200 p.i. were representative time points of the early and late phases of chronic infection, respectively (manuscript in preparation).

In this study, we evaluated an immunotherapy administrated during the early chronic phase of experimental *T. cruzi* infection. We hypothesized that a low dose BNZ treatment in combination with a therapeutic vaccine (TSA-1-C4 and Tc24-C4 recombinant antigens in a formulation with a synthetic TLR-4 agonist-adjuvant, E6020-SE) could provide a strong antigen-specific CD8^+^ T cell immunity, improving memory response and preventing the development of cardiac fibrosis.

This study could support the use of the vaccine-linked chemotherapy approach given in early chronic infection, preventing or delaying the development of severe manifestations in prolonged stages of CD.

## Materials and methods

### Ethics statements

All studies were approved by the institutional bioethics committee of the “Centro de Investigaciones Regionales Dr. Hideyo Noguchi”, Universidad Autónoma de Yucatán (Reference #CEI-08-2019) and were performed in strict compliance with NOM-062-ZOO-1999.

### Proteins, adjuvant and benznidazole

The recombinant TSA-1-C4 and Tc24-C4 antigens were obtained from the Centro de Investigación y Estudios Avanzados (CINVESTAV) of the Instituto Politécnico Nacional (IPN), Mexico. Each TSA-1-C4 or Tc24-C4 coding sequence was cloned into a pET41a+ *E. coli* expression vector. The resulting plasmid DNA was transformed into BL21 (DE3) cells induced with isopropyl-beta-D-1-thiogalactoside (IPTG) for protein expression. Recombinant proteins were purified by ion exchange (IEX) and size exclusion chromatography (SEC) (28, 30). The integrity and size of each recombinant protein were analysed by SDS-PAGE electrophoresis (**S1 Fig**). The recombinant proteins were formulated with the adjuvant E6020 in a stable squalene emulsion (SE). E6020 Toll-like receptor 4 agonist was acquired through Eisai, Inc (43). Benznidazole (N-Benzyl-2-nitro-1H-imidazole-1-acetamide) was obtained through Sigma Aldrich®; for its use, it was solubilized in 5% dimethyl sulfoxide (DMSO)-95% deionized water (32, 42).

### Mice and parasites

Female BALB/c mice (BALB/cAnNHsd) were obtained at 3-4 weeks old from ENVIGO-Mexico. Animals were housed in groups of 5 per cage, received *ad libitum* food and water and a 12-h light/dark cycle. Mice were acclimated for two weeks before starting the study. *T. cruzi* H1 parasites, originally isolated from a human case in Yucatan, Mexico were maintained by serial passage in BALB/c mice every 25 to 28 days and used for infections as previously described (44).

### Macrophages cell line

RAW 264.7 cell line was acquired from American type culture collection (ATCC® TIB-71™). Cells were cultured in DMEM medium (Gibco ®) supplemented with 10% fetal bovine serum (FBS, Gibco ®) and 1% penicillin/streptomycin (Gibco ®), in an atmosphere of 5% CO_2_ and 95% humidity at 37 °C. Cells were passaged in T-75 culture flasks (Corning ®) after reaching 80% confluence and were detached using 0.25% trypsin (Corning ®).

### Experimental infection and therapeutic treatment

A total of 70 mice were infected with 500 trypomastigotes of *T. cruzi* (H1 strain) by intraperitoneal injection. In order to confirm the infection, parasitemia was measured in Neubauer chamber by examination of fresh blood collected from the mouse tail at day 27 post-infection. (**S1 Table**). Survival was monitored up to day 200 post-infection (p.i) (**S2 Fig**). At day 72 p.i. (early chronic phase) surviving mice were randomly divided into groups of 8 individuals, the therapeutic vaccine was injected intramuscularly, and a vaccine boost with the same formulation was administrated one week after (day 79 p.i.). Each vaccine dose consisted of 12.5 μg of recombinant TSA-1-C4, 12.5 μg of recombinant Tc24-C4 and 5 μg of E6020-SE (28, 32). From day 72 to 79 p.i., mice were given daily 25 mg/kg BNZ by oral gavage, which corresponds to a 4-fold reduction in the conventional regimen of BNZ treatment. Additional groups of mice received the therapeutic vaccine alone (12.5 μg of TSA-1-C4, 12.5 μg of Tc24-C4 and 5 μg of E6020-SE), low dose BNZ alone (25 mg/kg), E6020-SE alone (5 μg) or saline solution as control. Groups that did not receive BNZ were given the vehicle solution (5% DMSO in 95% deionized water) by oral gavage. One additional control group with four non-infected mice was also included. At day 200 p.i., mice were euthanized using ketamine/xylazine-induced deep anaesthesia followed by cervical dislocation, and spleens and hearts were collected for further analysis (**Fig 1**).

**Fig 1.**
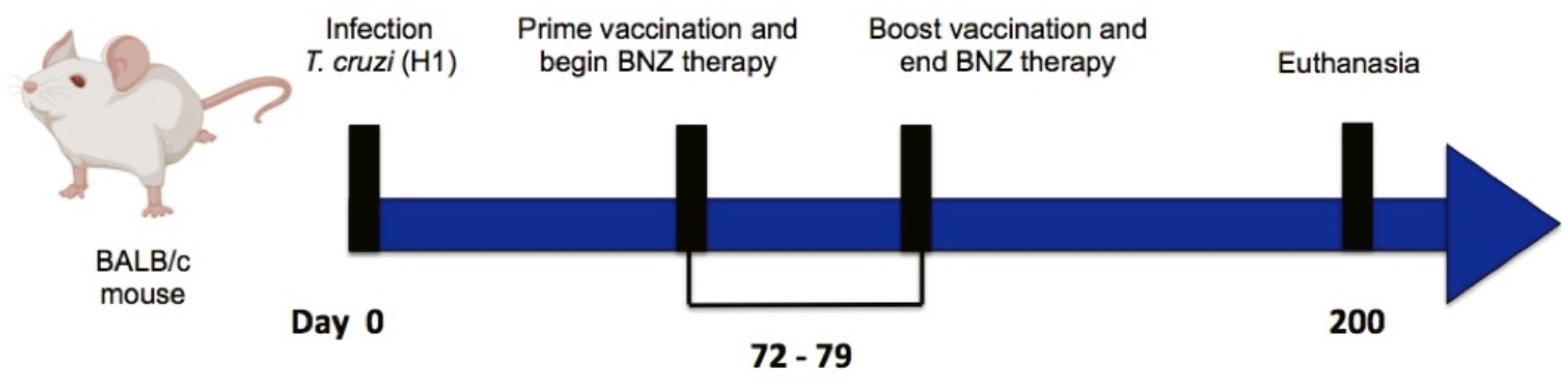
Timeline for experimental infection, prime-boost vaccination and euthanize using murine model.

### Quantification of parasites burden

Total DNA was isolated from cardiac tissue using the Kit Wizard® Genomic DNA purification (Promega Madison WI). Each sample was quantified employing a BioSpec-nano spectrophotometer system (SHIMADZU®) and adjusted to 50 ng of DNA from cardiac tissue, then quantitative real-time PCR (qPCR) was performed in an Illumina® EcoTM thermocycler using SYBR Green Master Mix® 1X and primers TCZ-F 5’-GCTCTTGCCCACAMGGGTGC-3’ and TCZ-R 5’-CCAAGCAGCGGATAGTTCAGG-3, which amplify a 188 pb product from the satellite region of *T. cruzi* DNA (45, 46). Cardiac parasite burdens were calculated based on a standard curve and expressed as parasite equivalents per 50 ng *T. cruzi*-DNA (47).

### Cardiac fibrosis and inflammation

For histopathological analysis, cardiac tissue from euthanized mice was fixed in 10% neutral buffered formalin solution, embedded in paraffin, cut into 5 μm sections, and stained with either Masson’s trichrome or haematoxylin and eosin (H&E) for fibrosis or inflammation infiltrate measurement respectively. To assess cardiac fibrosis or cardiac inflammation, five representative pictures from each slide stained were acquired at 10X magnification using an OLYMPUS microscope (CX23) adapted with a digital camera MiniVID P/N TP605100 (LW Scientific ®). Image analysis was performed using ImageJ software version 2.0.0/1.52p. To quantify cardiac fibrosis, pixels corresponding to fibrosis (blue coloured) were quantified and normalized by subtracting the average data obtained from non-infected control group to total pixels of the sample to assess the percentage of fibrotic area in the cardiac tissue (16,42,48). To quantify inflammatory infiltrate, the number of pixels corresponding to total nuclei was quantified and normalized to total cardiac tissue area (48, 49). Data is presented as cardiac fibrosis percentage area or cardiac inflammatory cells per mm^2^.

### Preparation of spleen mononuclear cells

Spleens were mechanically dissociated by being pressed through a 100 μm pore-size cell strainer. Splenocytes were rinsed with RPMI medium (Gibco ®) supplemented with 10% FBS and 1% penicillin-streptomycin (RPMIc) and pelleted by centrifugation for 5 min at 400 x *g* at room temperature. The supernatant was decanted, and the splenocyte pellet was resuspended in balanced salt solution buffer (BSS) pH 7.4. Afterwards, the splenocyte suspension was mixed with Ficoll-histopaque (GE Healthcare BIO-Sciences®) solution in 3:4 proportion and centrifuged at 400 x *g* for 40 min. The mononuclear cell layer was collected and washed twice with BSS buffer. The cell pellet was resuspended in RPMIc medium, cell viability was assessed by Trypan blue exclusion test and cell numbers were determined in a Neubauer chamber.

### Intracellular cytokine and memory T cell immune phenotyping

A total of 5×10^5^ mononuclear cells were co-cultivated with RAW 264.7 macrophages previously stimulated with TSA-1-C4+Tc24-C4 (25 μg/mL final concentration) in 10:1 proportion. Co-cultures were incubated in 5% CO_2_ and 95% humidity at 37°C during 20 h for intracellular cytokine production or 96 h for memory immune-phenotyping assays. To evaluate intracellular cytokine production, brefeldin A (BD biosciences ®) was added to co-culture for the last 4 hours of incubation. Re-stimulated cells were collected and washed twice with PBS+BSA 0.01%, then, cells were stained with anti-CD3 Alexa-647 (BD biosciences ®), anti-CD4 PE-Cy7 (BD biosciences ®) and anti-CD8 PERCP-Cy5.5 (BD biosciences ®) (30) or anti-CD3 PE-Cy7 (BD biosciences®), anti-CD4 APC (BD biosciences ®), anti-CD8 BB515 (BD biosciences ®), anti-CD44 PE (BD biosciences ®) and anti-CD62L BV510 (BD biosciences ®) for memory immune-phenotyping assay. For intracellular cytokine production, cells were fixed with Cytofix/Cytoperm (BD biosciences ®), and permeabilized according to the manufacturer’s instructions. Permeabilized cells were stained with anti-IFNγ (BD biosciences ®) and anti-Perforin (INVITROGEN ®). Cells were resuspended in FACS buffer and acquired on a BD FACSVerse flow cytometer. At least 50,000 total events in the mononuclear cell gate were obtained using FACSuite™ software version 1.0.5. Data were analysed in FlowJo software version 10.0.7r2. Stimulation index was calculated with the frequency of cells (stimulated with TSA-1-C4+Tc24-C4) and the frequency of non-stimulated cells (RPMIc alone). For intracellular analysis, stimulation index was measured with the median fluorescent intensity (MFI) of stimulated cells and the MFI of non-stimulated cells. A stimulation index > 1, indicates the presence of antigen-specific cells. Flow cytometry gating strategies for IFN*γ* and perforin expression or memory responses are presented in **S3 Fig** and **S4 Fig**.

### Statistical analysis

All tests were run in GraphPad Prism software version 6.0.c. Data were analysed by one-way ANOVA or Kruskal–Wallis tests for multiple groups, depending on its distribution followed by Tukey or Dunn’s *post hoc* test. When only two comparison groups were analysed, Student’s t-test or Mann–Whitney U-test was performed depending on data distribution. Differences between groups were considered statistically significant when *P*-value was less than 0.05.

## Results

### Vaccine-linked chemotherapy administered during the early chronic infection prevents cardiac fibrosis caused by *T. cruzi*

To evaluate the therapeutic efficacy of the vaccine-linked treatment, we measured cardiac *T. cruzi*-parasite burden, fibrosis, and inflammation. At day 200 p.i. *T. cruzi* cardiac parasite burden from mice treated and untreated was below the limit of detection of our qPCR test (<1 parasite per 50 ng of cardiac tissue) (**Fig 2A**). There were no differences comparing all experimental groups (Kruskal-Wallis, *P*=0.054). These data suggest that the methodology used has limitations in determining the therapeutic efficacy upon burden parasite of the formulation during the late chronic phase of infection in BALB/c mice.

**Fig 2.**
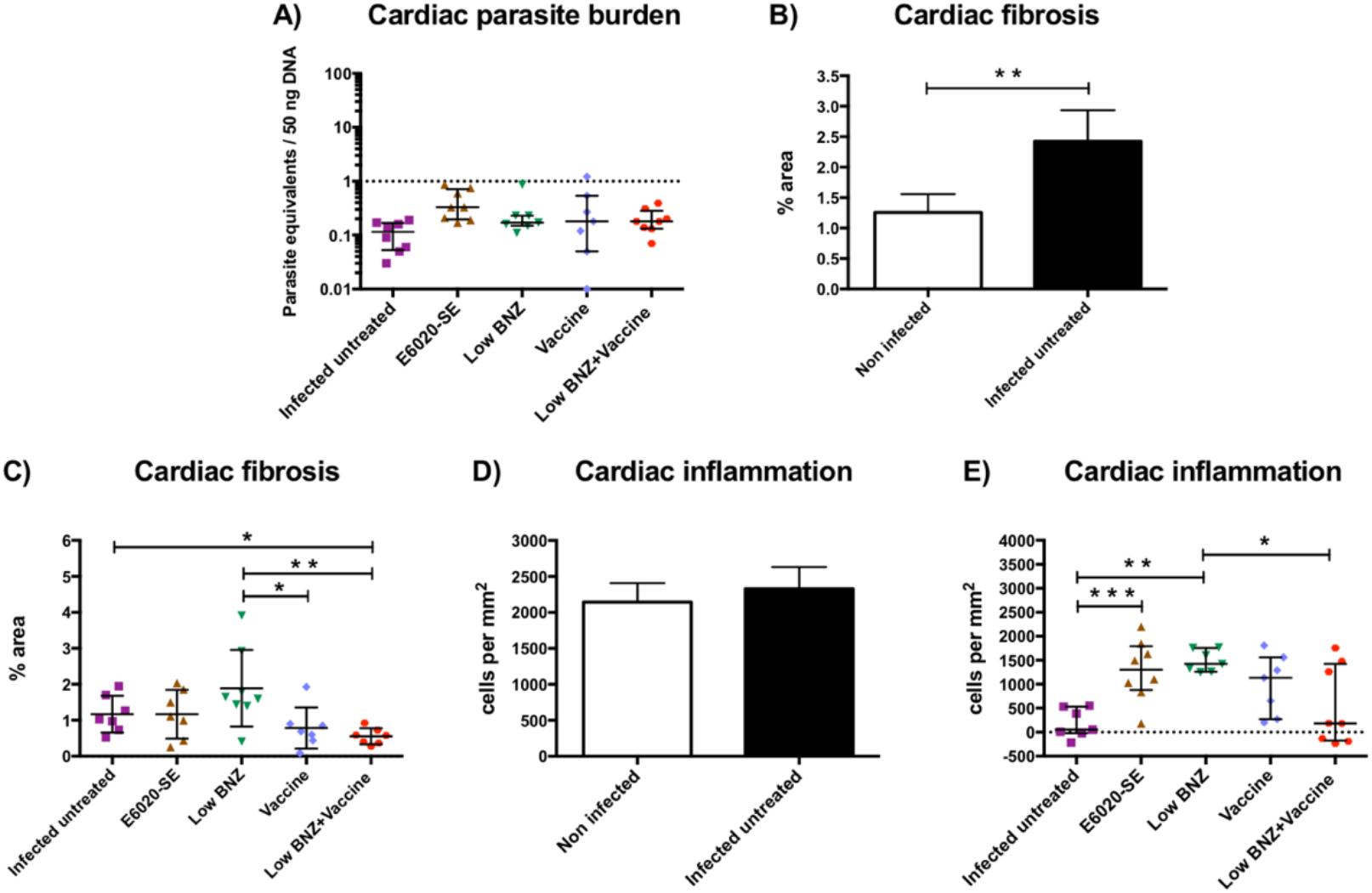
Protective effect of the vaccine-linked chemotherapy. Mice were euthanized at day 200 p.i and heart samples were collected. (**A**) Cardiac parasite burdens were quantified by quantitative real-time PCR. The dotted line represents the cut-off for the limit of detectable quantification (LOQ) based on serially diluted *T. cruzi*-enriched cardiac tissue DNA (1 parasite equivalents per 50 ng of DNA). Each point represents an individual mouse; horizontal lines denote median ± interquartile ranges values; significance calculated by Kruskal-Wallis test with Dunn’s correction for multiple comparisons. (**B**) Percentage fibrosis area in non-infected and infected untreated mice. Bars denote means and standard deviation; significance calculated by Student’s t-test. (**C**) Percentage fibrosis area for all experimental groups. The cardiac fibrosis was quantified from representative images of Massońs trichrome-stained tissue sections using Image J software. Each point represents an individual mouse, horizontal lines denote means ± SD; significance was calculated by Student’s t-test. (**D**) Infiltrate cells/mm^2^ in non-infected and infected untreated mice. Bars denote means and standard deviation; significance calculated by Student’s t-test. (**E**) Infiltrate cells/mm^2^ for all experimental groups. Cardiac inflammation was quantified from representative images of H&E-stained tissue sections using Image J software. Each point represents an individual mouse, horizontal lines denote median ± interquartile ranges values; significance was calculated by Mann-Whitney U-test. Significance is indicated as follows *, *P*≤0.05; **, *P*≤0.01; ***, *P*≤0.001.

On the other hand, we evaluated cardiac fibrosis in heart tissue sections collected from *T. cruzi*-infected mice at day 200 p.i. Representative images of Massońs trichrome stained-cardiac tissue from each experimental group are shown in **Fig 3A-F**. As we observed in **Fig 2B**, there was a significantly higher percentage of cardiac fibrosis in infected untreated mice (2.426 ± 0.51) compared to non-infected mice (1.259 ± 0.299) (Student’s t-test, *P=*0.003). Thus, mice infected with *T. cruzi* developed cardiac fibrosis as a consequence of chronic infection. Also, we found significant differences comparing the combination of low BNZ + vaccine (0.551 ± 0.223) with infected untreated mice (1.167 ± 0.510) or low BNZ alone treated mice (1.889 ± 1.065) (Student’s t-test, *P=*0.012 and *P=*0.006 respectively) (**Fig 2C**). This finding suggests that the vaccine-linked chemotherapy administered at early chronic infection prevents cardiac fibrosis caused by *T. cruzi* chronic infection. On the other hand, we observed significantly lower fibrosis percentage in vaccine alone treated mice (0.784 ± 0.573) compared to low BNZ alone group (Student’s t-test, *P=*0.029), suggesting that the use of the vaccine (TSA-1-C4+Tc24-C4+E6020-SE) is better than BNZ treatment at lower dose in our experimental infection model (**Fig 2C**).

**Fig 3.**
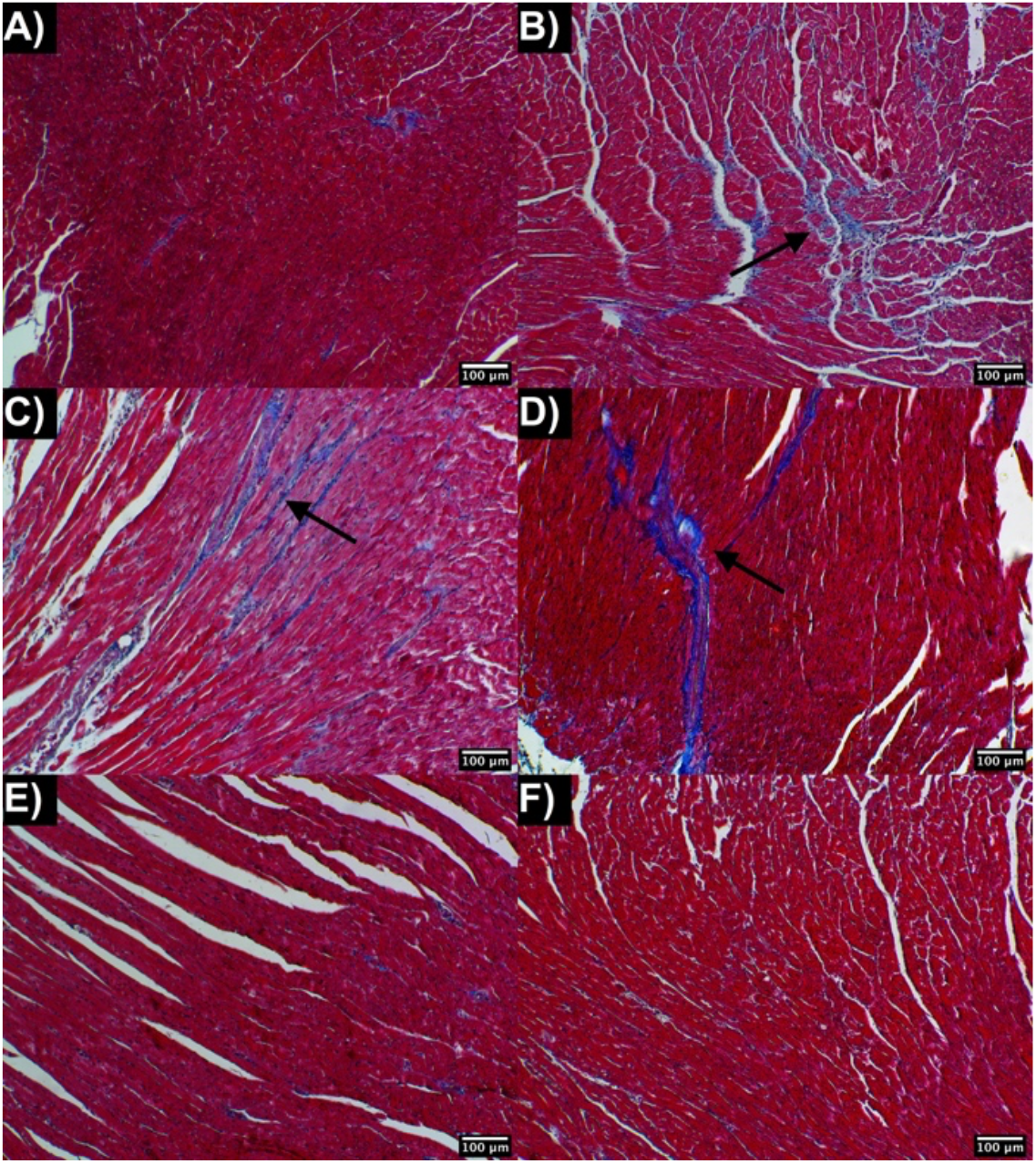
Representative images of Massońs trichrome stained-cardiac tissue from (**A**) non-infected, (**B**) infected untreated, (**C**) E6020-SE, (**D**) low BNZ, (**E**) Vaccine and (**F**) low BNZ plus vaccine experimental groups. Cardiac muscle appears in red and collagen fibbers in blue. Black arrows show collagen staining.

We also evaluated the inflammatory infiltrate in heart tissue sections collected from *T. cruzi*-infected mice at day 200 p.i. Representative images of H&E stained-cardiac tissue from the different experimental groups are shown in **Fig 4A-F**. According to **Fig 2D** we observed similar levels of inflammatory cell density between non-infected mice (2,143 ± 265.1) and infected untreated mice (2,329 ± 302.9), (Student’s t-test, *P=*0.335). Hence, our findings suggest that infected mice at late chronic infection (200 days p.i.) present basal levels of infiltrating inflammatory cells. Moreover, a significantly lower proportion of infiltrating inflammatory cells was found comparing infected untreated mice with E6020-SE alone (1290 ± 636.4) or low BNZ alone (1485 ± 223.7) groups (Mann-Whitney U-test, *P=*0.002 and P<0.001 respectively) (**Fig 2E**), suggesting that, using this experimental infection model, either adjuvant or BNZ treatments given alone contributes to the development of cardiac inflammation. Interestingly, we observed that the low BNZ + vaccine-treated group (537.5 ± 820.9) had significantly lower levels of inflammatory cell density compared to low dose BNZ-treated group (Mann-Whitney U-test, *P=*0.04) (**Fig 2E**), indicating that there is a benefit adding the vaccine formulation to the BNZ alone treatment.

**Fig 4.**
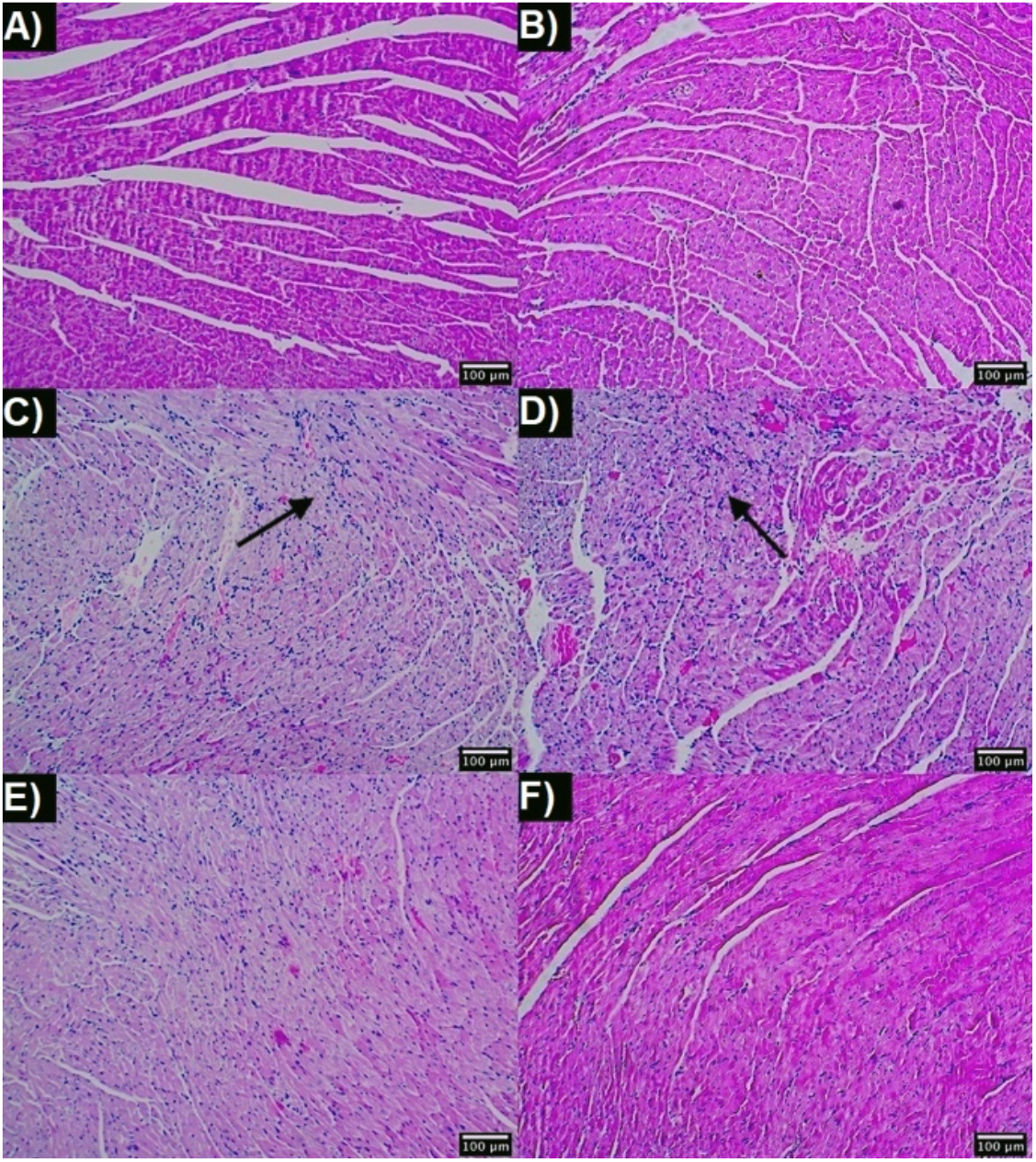
Representative images of H&E stained-cardiac tissue from (**A**) non-infected, (**B**) infected untreated, (**C**) E6020-SE, (**D**) low BNZ, (**E**) Vaccine and (**F**) low BNZ plus vaccine experimental groups. Cardiac muscle appears in pink and the nuclei of infiltrating inflammatory cells appear in purple. Black arrows show the presence of infiltrating inflammatory cells.

### Immunotherapy with low BNZ plus vaccine primes a cytotoxic profile in CD8^+^ and CD4^+^ T antigen-specific cells

We evaluated the presence of antigen-specific CD4^+^ and CD8^+^ T cells in mice during the late chronic infection. As shown in **S5 Fig**, all experimental groups had a mean of stimulation index ≤ 1 for CD4^+^ T cells population; suggesting that CD4^+^ T cells from infected mice (regardless of treatment) have no detectable TSA-1-C4+Tc24-C4-antigen specific cells. Besides, the stimulation index of antigen-specific CD8^+^ T cells from all groups had a mean of stimulation index ≥ 1 (**S5 Fig**), confirming the presence of TSA-1-C4+Tc24-C4-antigen specific cells in *T. cruzi*-infected mice at the late chronic phase. These results suggest that during the late chronic phase there are antigen specific CD8^+^ T cells in all infected groups that can be recalled using antigen presenting cells (APC) such as macrophages. However, at 200 days p.i., we did not find differences among groups for antigen-specific CD4^+^ or CD8^+^ T cells (ANOVA, *P=*0.858 and *P=*0.793 respectively).

To evaluate the intracellular IFNγ and perforin production by TSA-1-C4+Tc24-C4- antigen-specific CD4^+^ and CD8^+^ T cells, mononuclear cells were isolated and co-cultivated with antigen-specific RAW 264.7 macrophages, as described before. According to **Fig 5A**, we observed that CD4^+^ T cells from the low BNZ + vaccine treated mice showed a significantly higher production of IFNγ (1.056 ± 0.062), compared to E6020-SE (0.958 ± 0.023) or low BNZ alone (0.954 ± 0.06) treated groups (Student’s t-test, *P=*0.002 and *P=*0.007 respectively). On the other hand, mice treated with the vaccine alone (1.367 ± 0.504) or the combination low BNZ + vaccine (1.089 ± 0.572) had the highest production of perforin by antigen-specific CD4^+^ T cells (**Fig 5B**) and were significantly higher when comparing with infected untreated mice (0.532 ± 0.359) (Mann-Whitney U test, *P=*0.002 and *P=*0.049 respectively).

**Fig 5.**
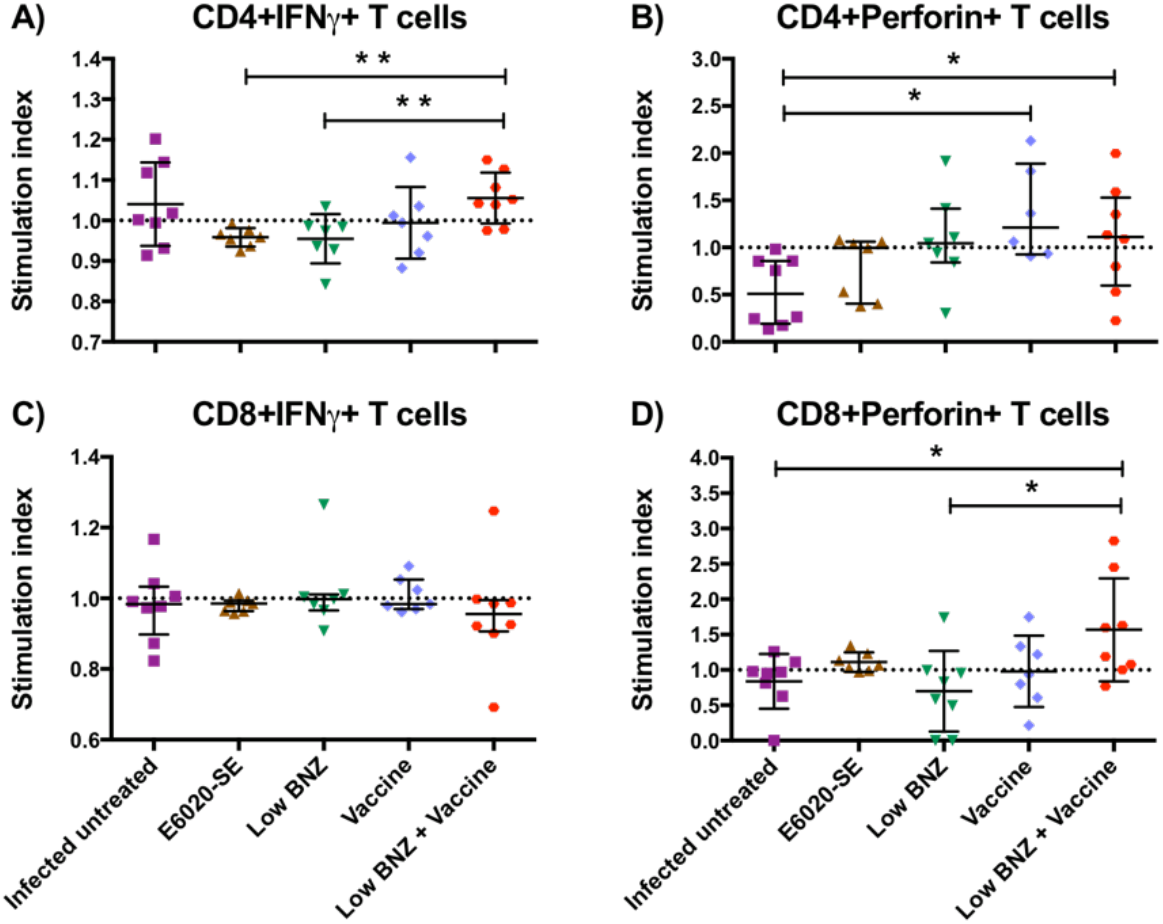
Effect of vaccine-linked chemotherapy on antigen-specific CD4^+^ and CD8^+^ T cells functional profile. Mononuclear cells isolated from mice at 200 days p.i were co-cultivated with macrophages stimulated *in vitro* with TSA-1-C4+Tc24-C4 (25 μg/mL) for 20 h. Data were analysed using FlowJo X software. Stimulation index of (**A**) antigen-specific IFNγ-producing CD4^+^ cells, (**B**) perforin-producing CD4^+^ cells, (**C**) IFNγ-producing CD8^+^ cells and (**D**) perforin-producing CD8^+^ cells are presented. Each point represents an individual mouse, horizontal lines denote means ± SD or median ± interquartile ranges values according to the normality of the data. Data were analysed using Student’s t-test or Mann-Whitney U-test. Significance is indicated as follows *, *P*≤0.05; **, *P*≤0.01.

For antigen-specific CD8^+^ T cells, we observed a cytotoxic immuno-phenotype profile characterized by higher perforin-producing CD8^+^ T cells in low BNZ + vaccine treated mice (**Fig 5D**), (1.567 ± 0.728) compared to the infected untreated group (0.838 ± 0.386) or the low BNZ treated group (0.699 ± 0.570) (Student’s t-test, *P=*0.025 and *P=*0.018 respectively), however, no differences were found when we evaluate the production of IFNγ by CD8^+^ T cells (**Fig 5C**). All these findings suggest that the vaccine-linked chemotherapy might ameliorate the fibrosis in *T. cruzi* chronic infection by induction of a TSA-1-C4+Tc24-C4- antigen specific CD4^+^ and CD8^+^ cytotoxic T cells with a perforin-phenotype and these can be recalled up to day 200 p.i. in our mouse model.

### Treatment given during early chronic infection induced a long-lasting *T. cruzi*-immunity

With the purpose to evaluate the memory T cell profile induced by the vaccine-linked chemotherapy, we measured markers of central (T_CM_) (CD44^+^CD62L^+^) and effector (T_EM_) (CD44^+^CD62L^-^) T cell memory subpopulation in CD4^+^ and CD8^+^ T cells.

As observed in **Fig 6A**, we found a significantly higher stimulation index of antigen-specific CD4^+^ T_CM_ sub-population in the low BNZ + vaccine treated mice (1.19 ± 0.073) and vaccine alone groups (1.206 ± 0.105) compared to infected untreated mice (1.085 ± 0.07) (Student’s t-test, *P=*0.026 and *P=*0.018 respectively). Similarly, a significantly higher stimulation index of CD8^+^ T_CM_ sub-population (**Fig 6B**), was observed in low BNZ + vaccine treated mice (1.485 ± 0.519) and vaccine alone groups (1.33 ± 0.18) compared to infected untreated mice (0.995 ± 0.149) (Student’s t-test, *P=*0.032 and *P=*0.002 respectively). Otherwise, we observed that low BNZ + vaccine treated mice (0.808± 0.054) and low BNZ alone treatment (0.808 ± 0.07) had lower stimulation index of antigen-specific CD4^+^ T_EM_ sub-population compared to infected untreated mice (0.908 ± 0.076) (Student’s t-test, *P=*0.012 and *P=*0.02 respectively) **Fig 6C**. Finally, there was no difference in CD8^+^ T_EM_ at late chronic infection (200 days p.i.) (**Fig6 D**). Of note, the vaccine alone or linked-chemotherapy elicited TSA-1-C4+Tc24-C4 antigen-specific central memory response during the late chronic phase of *T. cruzi*-infection in BALB/c mice.

**Fig 6.**
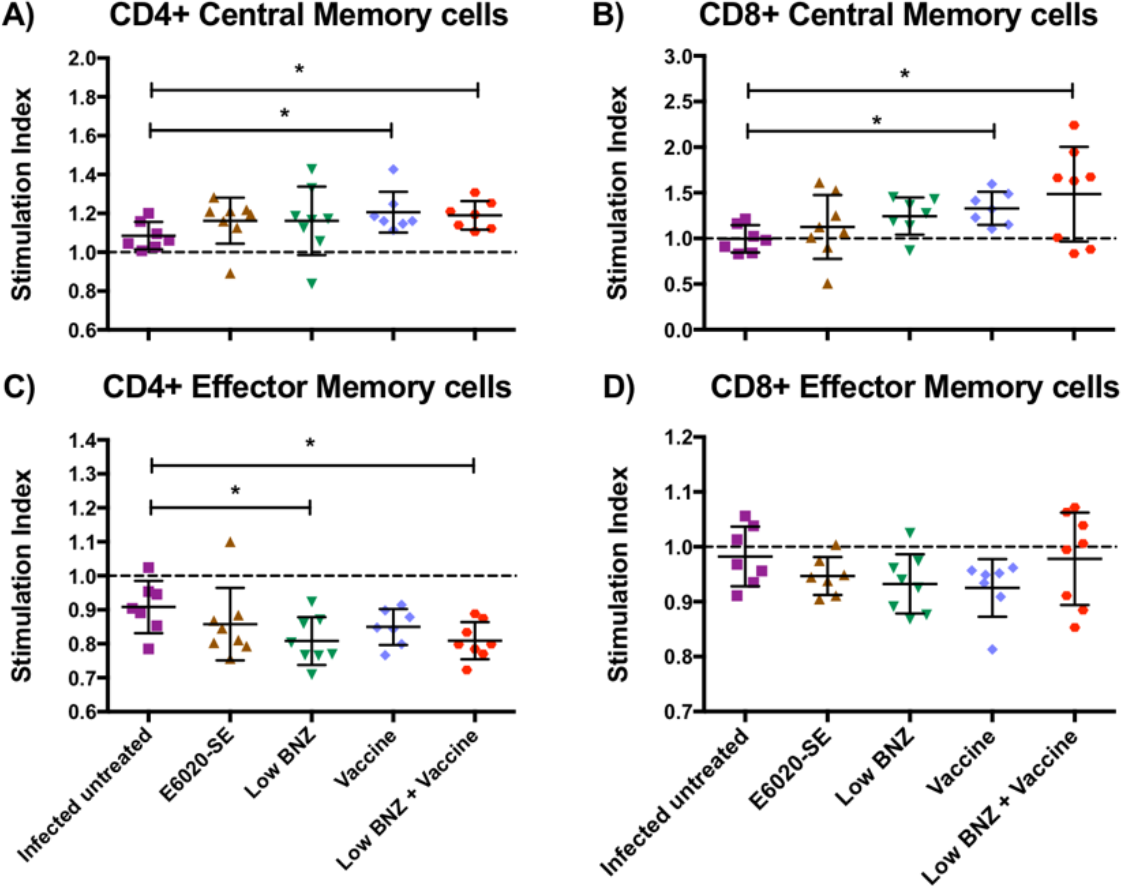
Effect of vaccine-linked chemotherapy on antigen-specific CD4^+^ and CD8^+^ T cells memory response. Mononuclear cells isolated from mice at 200 days p.i. were co-cultivated with macrophages stimulated *in vitro* with TSA-1-C4+Tc24-C4 (25 μg/mL) for 96 h. Data were analysed using FlowJo X software. Stimulation index of antigen-specific (**A**) CD4^+^ Central Memory, (**B**) CD4^+^ Effector Memory, (**C**) CD8^+^ Central Memory and (**D**) CD8^+^ Effector Memory T cells are presented. Each point represents an individual mouse, horizontal lines denote mean ± SD values, and data were analysed using Student’s t-test. Significance is indicated as follows *, *P*≤0.05.

## Discussion

One of the greatest challenges in chronic CD is the development of therapies that improve prognosis and, even, reverse the cardiac injury. Our research partnership is developing a therapeutic vaccine against CD that seeks to prevent or delay the onset of CCC in infected patients. Previous studies by our group have evidenced the feasibility of DNA vaccines formulated with TSA-1 and Tc24 *T. cruzi*-antigens in mice and dogs (33,50,51). This DNA-bivalent vaccine was associated with a CD8^+^ T cell activity, IFNγ production, Th1 immune response. Since DNA vaccines historically have not been yet progressed successfully to emergency use authorization or full licensure for use in humans, our studies have focused on recombinant protein antigens (52). Hence, we embarked on studies to evaluate vaccines formulated with TSA-1 or Tc24 recombinant proteins in conjunction with Toll-like receptor 4 agonist adjuvants (30, 47). These previous studies have concluded that either Th1 or balanced Th1/Th2 immune responses are associated with reductions in parasite burdens, fibrosis, and host infiltrating inflammation linking it to confer protection in experimental models of acute *T. cruzi* infection (16,31,32).

Specific genetic mutations have been made for both recombinant proteins, to facilitate production and increase stability, while maintaining immunogenicity in mice; the proteins resulting were named as TSA-1-C4 and Tc24-C4 (28, 29). These designations reflect the modification of cysteines to prevent protein aggregation from intermolecular disulphide bond formation. The bivalent vaccine, therefore, is comprised of two recombinant *T. cruzi* antigens, Tc24-C4 and TSA-1-C4, demonstrated immunogenicity in non-human primate trials study (53). An effective therapeutic treatment for CD must prove its effectiveness at preventing or delaying the onset of CCC (17). Thereby, it is critical to evaluate therapeutic treatments in pre-clinical models during the chronic phase of *T. cruzi*-infection. Recently, studies aimed to evaluate BNZ in reduced dosing regimens to decreasing adverse side-effects (39, 41). Here, we evaluated the therapeutic efficacy of a vaccine candidate formulated with TSA-1-C4 + Tc24-C4 recombinant antigens combined with a 4-fold reduction in the amount and dosage regimen of BNZ treatment given during the early chronic phase of *T. cruzi*-infection in a murine model and followed until the late chronic phase of the disease.

In terms of protective effect, a diminished parasitemia and cardiac parasite burdens are related with therapeutic efficacy in murine models of *T. cruzi* acute infection (15,31,32,42,47). On the other hand, *T. cruzi* parasites in blood and cardiac tissue decrease with time during *T. cruzi* chronic infection becoming undetectable in blood and restricted in tissues, where they are not always demonstrable, even by using highly sensitive amplification techniques as qPCR assay (54). However, novel techniques using a bioluminescence imaging system have allowed to measure the burden parasite *in vivo* during *T. cruzi*-chronic infection demonstrating that the *T. cruzi* presence in the heart is spatially dynamic (55). This finding suggests that parasitemia, as well as cardiac parasite burden alone, are not robust indicators to evaluate therapeutic efficacy during *T. cruzi* chronic infection in murine models. Therefore, the absence of detectable parasites in the heart at a point of infection does not necessarily exclude the ongoing infection itself, as the parasite could be restricted to other organs, such as in the gastrointestinal tract (55). This could explain our results, as we detected low levels of parasite burden in the heart. Further studies need to be performed in order to understand the mechanism that allows the establishment of cardiac disease in irregular levels of parasites.

Beyond parasite reduction, there are findings stressing the role of reducing both cardiac fibrosis and inflammation in patients, non-human primates and mice (56, 57). In fact, the use of non-invasive methods to measure fibrosis have allowed to distinguish potentially useful biomarkers of cardiac fibrosis, such as TGF-*β*, connective tissue growth factor, and platelet-derived growth factor-D (48). In the current study, we showed that infected untreated mice in chronic infection exhibited a more severe cardiac fibrosis compared to the non-infected control group, as expected. Also, we observed that the vaccine-linked chemotherapy-treated group had on average only 50% of the cardiac fibrosis area compared to infected untreated group. Thus, we demonstrated that the vaccine-linked chemotherapy given at the early chronic phase of *T. cruzi* infection is able to prevent cardiac fibrosis in our mouse model. In this experimental infection model, the administration of the low dose of BNZ alone (25mg/kg for 7 days) is unable to prevent cardiac fibrosis at 200 days p.i. These results coincide with previous studies, showing that BNZ therapy (100mg/kg for 20 days) prevents the development of cardiac fibrosis in the murine model when treatment is administrated in the acute phase however, the drug fails when is administrated during the chronic phase of infection (58).

Our findings support previous studies in mice during acute infection by *T. cruzi* showing that Tc24 and Tc24-C4 immunizations or the vaccine-linked chemotherapy can reduce parasite burden, cardiac fibrosis, and inflammation (15,16,32,42). In addition, we confirmed that the protective effect is provide for the antigens and it is not due to adjuvant. For instance, we have previously evaluated the dose-response effect of E6020-SE administration on cardiac fibrosis using a lethal acute infection murine model (59). According to our previous findings, E6020-SE administration alone does not prevent cardiac fibrosis development during acute phase. Here, we showed that mice treated with 5 μg of E6020-SE alone during the early chronic infection, still develop cardiac fibrosis at the late chronic phase of *T. cruzi*-infection. Therefore, our results suggest that the vaccine antigen component is necessary to achieve optimal protection against cardiac fibrosis development.

As part of the study, we evaluated protection against cardiac inflammation conferred by our vaccine-linked chemotherapy. According to our data, no significant differences were observed when we compared our vaccine-linked chemotherapy-treated group with the infected untreated group. However, we observed similar levels of inflammatory infiltrate in cardiac tissue from infected untreated and non-infected mice suggesting that the percentage of inflammation in cardiac tissue from infected mice decrease over the course of infection until reach basal levels on late chronic stages of the disease (200 days p.i), as has been previously described by Hoffman (48).

The CD8^+^ cytotoxic T cells activation is essential to achieve immunity against *T. cruzi* as well other parasites (60, 61). We showed that the vaccine-linked chemotherapy-treated group had on average double production of perforin by antigen-specific CD8^+^ T cells compared to infected untreated mice. Our results pointed out that the immunotherapy with TSA-1-C4+Tc24-C4 recombinant antigens is able to prime a cytotoxic profile (CD8^+^Perf^+^) during the late chronic phase of *T. cruzi*-infection. Also, vaccine-linked chemotherapy elicited an antigen-specific CD4^+^ CTL sub-population, these cells can secrete cytotoxic granules that directly kill infected cells in an MHC-class-II-restricted context. Previous studies have described CD4^+^ CTL sub-population in both, human and murine models (62–64). Currently, the mechanism used by CD4^+^ CTL is unclear, however; this sub-population can exhibit functions complementary to CD8^+^ CTLs (65). More studies are needed to elucidate the role of CD4^+^ CTL in *T. cruzi* infection.

On the other hand, we did not observe significant differences neither antigen-specific CD4^+^IFNγ^+^ nor CD8^+^IFNγ^+^-producing T cells in the vaccine-linked chemotherapy-treated mice compared with the infected untreated group. Both phenotypes are characteristic of the Th1 immune response that is known to confer protection against *T. cruzi* acute infection. We believe that this finding is consistent with the chronic phase of infection since parasitemia has been controlled as well as cardiac parasite burden in both infected mice groups. Therefore, it is unlikely to observe the activation of TSA-1-C4 or Tc24-C4 antigen-specific CD4^+^IFNγ^+^ or CD8^+^IFNγ^+^ T cells populations in the spleen of mice that have already controlled the infection.

A challenge in the development of an effective vaccine against *T. cruzi* is the induction of long-lived memory cells, which confers long-term protection. In our study, we were able to recall antigen-specific CD4^+^ and CD8^+^ T_CM_ sub-populations at 200 days p.i. The central memory T cells are distinguished for having a proliferative response followed by antigenic stimulation that live longer than effector memory cells. Bixby and Tarleton have previously reported during *T. cruzi*-infection in mice this sub-population in CD8^+^ T cells showing distinctive features called as T_CM_ cells (36). Similarly, T_EM_ sub-population represents a type of terminally differentiated cells that produce IFN*γ* and IL-4 or contain prestored perforin (34). In this study, we observed a low proportion of either CD4^+^ or CD8^+^ T_EM_ sub-populations regardless of treatment from infected mice at 200 days p.i probably as consequence that effector cells are characterized by requiring a continuous antigen presentation by APC in order to proliferate and differentiate. We suggested that during the first weeks after the vaccine administration, T_EM_ sub-populations have differentiated and performed their effector activity, preventing the dissemination of the parasite, and ensuring the survival of the mice. This may explain the low levels of load parasite load observed by qPCR at day 200 p.i. Thus, we propose that for this experimental infection model, during the late chronic phase, the immune system remains in a “resting-state”, in which there is a limited effector activity. However, the use of the recombinant-protein vaccine recalls a strong central memory response mediated by antigen-specific T cells at 200 days p.i. In sum, our results suggest that a TSA-1-C4+Tc24-C4 antigen-specific T central memory sub-population could protect against reinfection during late chronic *T. cruzi* infection in our murine model.

There are some limitations in this study. We did not measure clinical parameters to assess heart function. Therefore, we were not able to correlate the reduced fibrosis with improved clinical cardiac output in our chronically infected mice. In addition, the BALB/c model of experimental infection used here seems to have an intrinsic resistance to *T. cruzi* acute infection allowing them to progress until the late chronic phase, which is characterized by lower levels of parasite burden and inflammation but higher percentages of cardiac fibrosis in infected untreated mice. However, our BALB/c model may be representative of a majority proportion of the *T. cruzi* infected human population that has the ability to control parasite burden and inflammation, remaining in asymptomatic chronic phase of CD for life.

## Conclusion

We demonstrate that treatment with a low dose BNZ and a vaccine immunotherapy protects mice against cardiac fibrosis progression and induces a long-lasting *T. cruzi*-immunity that persists for at least 18 weeks post-treatment. This study supports the use of a vaccine-linked chemotherapy approach given in early chronic infection, however; additional studies in other preclinical models that develop CCC and with more characteristics of human disease, such as non-human primates, will be necessary before the combination of a vaccine-linked to BNZ can be moved into clinical trials.

## Supporting information

**S1 Table. Parasitemia measurement.** A total of 70 BALB/c mice were infected with 500 trypomastigotes of *T. cruzi* (H1 strain) by intraperitoneal injection. A total of 4 mice were used as non-infected control group and received only saline solution. Parasitemia was measured in Neubauer chamber by examination of fresh blood collected from the mouse tail at day 27 post-infection. All mice who received infection were positive for *T. cruzi*, while neither mouse from non-infected control group were reported as positive. **SD**, standard deviation; **CI**, confidence interval.

**S1 Fig. protein integrity assessment.** SDS-PAGE analysis at 12% of acrylamide/bis-acrylamide and stain with PageBlue^TM^ Protein Staining Solution (ThermoFisher Scientific*®*). Molecular weight marker (Spectra^TM^ Multicolor Broad Range Protein Ladder from ThermoFisher Scientific*®*) and recombinant proteins TSA-1-C4 (65 kDA) and Tc24-C4 (24 kDa) are presented.

**S2 Fig. Survival curve.** A total of 70 BALB/c mice were infected with 500 trypomastigotes of *T. cruzi* (H1 strain) by intraperitoneal injection. A total of 4 mice were used as non-infected control group and received only saline solution. Survival was monitored during 200 days post-infection.

**S3 Fig. Flow-cytometry gating strategy for IFNγ and perforin production.** The dot-plots show the mononuclear cells gating based on (**A**) forward-scatter (FSC) and side-scatter (SSC) properties, (**B**) doublets exclusion, (**C**) identification of CD3^+^ positive cells, (**D**) phenotype of CD4^+^ and CD8^+^ cells and (**E**) IFN*γ* and perforin expression. Gates were stablished based on FMO controls corresponding to each antibody.

**S4 Fig. Flow-cytometry gating strategy for central and effector memory response.** The dot-plots show the mononuclear cells gating based on (**A**) forward-scatter (FSC) and side-scatter (SSC) properties, (**B**) doublets exclusion, (**C**) identification of CD3 positive cells, (**D**) phenotype of CD4^+^ and CD8^+^ cells and (**E**) central memory and effector memory profile defined by (CD44^+^CD62L^+^) and (CD44^+^CD62L^-^) expression respectively. Gates were established based on FMO controls corresponding to each antibody.

**S5 Fig. Effect of vaccine-linked chemotherapy on antigen-specific total CD4^+^ and CD8^+^ T cells response.** Mononuclear cells isolated from mice at 200 days post-infection were co-cultivated with macrophages stimulated in vitro with TSA-1-C4+Tc24-C4 (25 μg/mL) for 20 h. Data were analysed using FlowJo X software. Stimulation index of antigen-specific (**A**) CD4^+^ cells and (**B**) CD8^+^ cells are presented. Each point represents an individual mouse, data were analysed using ANOVA for multiple comparisons.

## Author contributions

**Conceptualization:** Dumonteil E, Cruz-Chan J.V and Villanueva-Lizama L.E

**Data curation:** Dzul-Huchim V.M and Villanueva-Lizama L.E

**Formal analysis:** Dzul-Huchim V.M, Cruz-Chan J.V and Villanueva-Lizama L.E

**Funding acquisition:** Rosado-Vallado M.E, Hotez P and Bottazzi M.E

**Investigation:** Dzul-Huchim V.M and Villanueva-Lizama L.E

**Methodology:** Dzul-Huchim V.M, Arana-Argaez V.E, Dumonteil E and Villanueva-Lizama L.E

**Project administration:** Hotez P, Bottazzi M.E and Villanueva-Lizama L.E

**Resources:** Ramirez-Sierra M.J, Martinez-Vega PB, Ortega-Lopez J and Gusovsky F

**Supervision:** Rosado-Vallado M.E, Cruz-Chan J.V, Villanueva-Lizama L.E, Hotez P and Bottazzi M.E

**Visualization:** Dzul-Huchim V.M and Villanueva-Lizama L.E

**Writing – original draft:** Dzul-Huchim V.M

**Writing – review & editing:** Arana-Argaez V.E, Cruz-Chan J.V, Villanueva-Lizama L.E, Ortega-Lopez J, Gusovsky F, Hotez P and Bottazzi M.E

